# Beyond Pairwise Interactions: Higher-Order Dynamics in Protein Interaction Networks

**DOI:** 10.1101/2022.05.03.490479

**Authors:** Kevin A. Murgas, Emil Saucan, Romeil Sandhu

## Abstract

Protein interactions form a complex dynamic system that shapes cell phenotype and function; in this regard, network analysis is a powerful tool for studying the dynamics of cellular processes. Graph-based models are limited, however, in that these models consider only pairwise relationships. Higher-order interactions are well-characterized in biology, including protein complex formation and feedback or feedforward loops. These higher-order relationships are better represented by a hypergraph as a generalized network model. Here, we present an approach to analyzing dynamic gene expression data using a hypergraph model and quantify network heterogeneity via Forman-Ricci curvature. We observe, on a global level, increased network curvature in pluripotent stem cells and cancer cells. Further, we use local curvature to conduct pathway analysis in a melanoma dataset, finding increased curvature in several oncogenic pathways and decreased curvature in tumor suppressor pathways. We compare this approach to a graph-based model and a differential gene expression approach.

## 1 Introduction

Cells are complex biological systems formed of thousands of interacting components, namely proteins and chemical molecules [1, 2]. The dynamic regulation of gene expression into specific protein levels and the numerous molecular interactions resulting among these proteins (i.e. the interactome) shape the phenotype of a cell, which manifests as a distinct cell type with specific functions. To study the factors that influence cellular phenotype, transcriptomic profiles can be experimentally measured via high-dimensional gene expression assays such as single-cell RNA-sequencing (scRNA-seq), allowing researchers to ask incisive biological questions of how cellular dynamics relate to physiological and pathological processes [3, 4]. For example, differential gene expression can help to identify driver genes or tumor suppressors in cancer [5]. Further, gene expression measurements can indicate activation or inactivation of molecular pathways that influence differentiation and cell type, as in stem cell differentiation or tumorigenesis. Modeling these cellular dynamics could ultimately inform therapeutic strategies (wound-healing, targeted cancer therapy, etc.) by predicting genes or pathways to target and manipulate the cell phenotype.

Classic differential gene expression typically considers genes independently, overlooking the relationships between genes or proteins within the cell [6, 7, 8]. In this regard, protein-protein interaction (PPI) network modeling provides an approach to investigate how the pattern of interactions between proteins contribute to cellular dynamics [9]. Protein interactions are determined by a number of experimental methods, including yeast-two-hybrid and co-precipitation protocols, and are compiled in curated databases such as KEGG and Reactome and integrative databases such as Pathway-Commons and STRINGdb, ultimately providing researchers with comprehensive protein interaction datasets to construct PPI network models [10, 11, 12, 13, 14, 15]. By formalizing the system of protein interactions as a network, one can begin to model how the pattern of interactions influence cellular phenotype and function, to then elucidate molecular mechanisms of healthy physiology and disease [15, 16, 17]. For example, one can examine the presence of redundancy in the network which may protect from failure in the case of individual component failure (i.e. gene mutation), or positive feedback loops which might lead to unstable behavior such as uncontrolled growth observed in cancers [18, 19, 20, 21].

Most current analyses of PPI networks, however, are limited in that the standard graph model considers only pairwise interactions (i.e. edges between two proteins at most) and not higher-order interactions [9, 22]. In biology, multiple proteins often work together with shared function; in fact, cells naturally exhibit protein complexes composed of several associated proteins [1, 23, 24] (Fig. 1A). Additionally, molecular pathways in cells can include signaling cascades involving multi-protein interactions, feedback or feedforward loops as well as cross-talk and overlap between pathways, but pairwise interactions in the graph model alone would only represent small segments of any given pathway of interest [11, 18, 19, 25, 26]. For these reasons, a generalized network model is necessary to effectively model higher-order relationships in biological networks.

**Figure 1:**
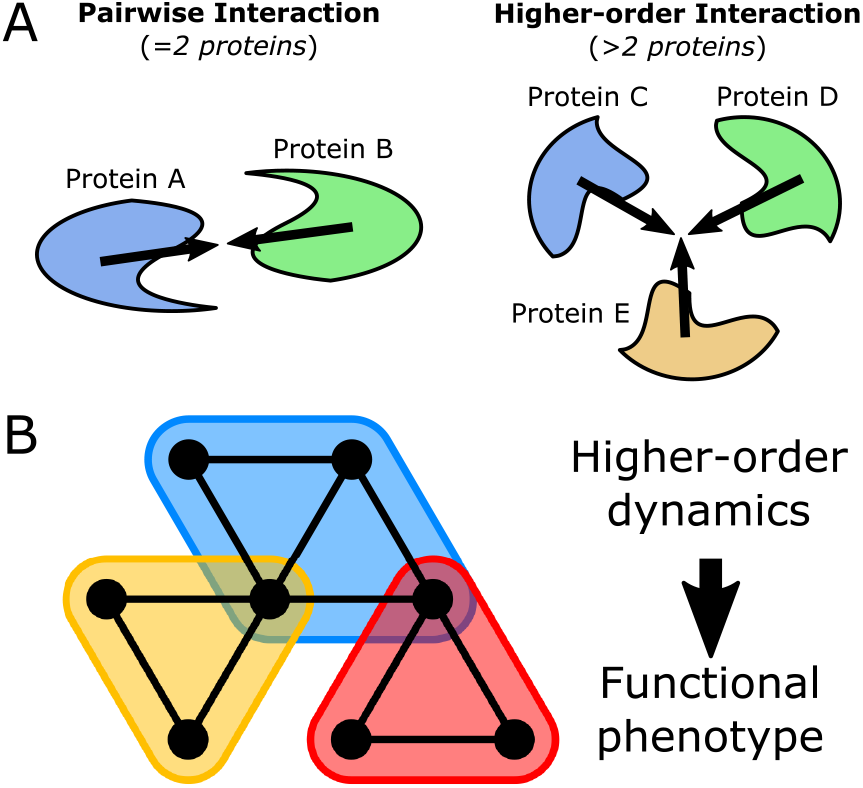
Hypergraph model of higher-order protein interactions. A: Illustration of two types of protein interactions: pairwise interaction between 2 proteins, and higher-order interaction between *>*2 proteins. B: Hypergraph model of higher-order interactions (shaded bubbles) among multiple proteins (black vertices).

This higher-order organization observed in protein interactions is best modeled as a hypergraph, a generalized network representation which considers higher-order relationships among multiple elements [27, 28, 29]. A hypergraph extends the standard graph model by considering relationships with any number of elements, generalizing the strictly pairwise edges of a graph. In the context of protein interactions, a “hyperedge” in a hypergraph model might contain any number of proteins to represent a single higher-order interaction. Further, there is a need not only to apply such higher-order models but, critically, to develop approaches which can use these models along with measured data (i.e. gene expression) to assess network dynamics and provide biological insight into cellular function and phenotype, with especial emphasis on identifying disease mechanisms and therapeutic targets.

In understanding dynamics, two primary concepts are equilibrium and heterogeneity [30, 31]. When a cancer expresses a certain pathway that drives growth and survival, how stable is this pathway to perturbation (i.e. targeted therapy)? This question is critical in understanding cancer drug response and how drug resistance arises [32, 33]. A heterogeneous interaction network with redundant, parallel communication might allow for the cancer to divert to alternative pathways to circumvent therapeutic targets [34, 35]. Heterogeneity of interaction dynamics is also fundamental in the process of cellular differentiation. Pluripotent cells, with the capacity to differentiate into several lineages, eventually reach points of bifurcation and commitment to express a unique set of proteins and molecular pathways that give the cell a specific functional phenotype [36, 37]. Understanding how these dynamics unfold is relevant to developing optimal therapeutic strategy, for example how to treat a cancer effectively by targeting its instability while also preventing drug resistance from developing [32, 33].

Recent studies have explored weighted PPI network models that incorporate geneexpression measurements as estimates of protein levels to calculate stochastic rates of interaction, allowing quantitative examination of how PPI network dynamics vary with gene expression in different biological settings [38, 39, 40]. Statistical properties that measure heterogeneity in a system, such as entropy and Ricci curvature, directly relate to the dynamic property of robustness, or stability under perturbation [39, 41]. These measures have been demonstrated to quantitatively indicate cellular pluripotency, or “stem-ness” of a cell, as well as cancer status and the increased robustness observed in cancer [31, 39, 42, 43]. While these previous studies utilized pairwise graph models, higher-order network models such as the hypergraph described above have yet to be explored. Importantly, a higher-order network model can be readily examined in terms of similar statistical properties to quantify heterogeneity and dynamic robustness [44].

In this study, we develop a hypergraph model of the protein interaction network based on a 2-dimensional simplicial complex, in which we extend the graph model to include 2-dimensional “faces” among subsets of vertices with shared edges (Fig. 1B,C) [44, 45, 46]. By considering a higher-dimensional network structure, we aim to account for higher-order interactions in the PPI network, including feedforward and feedback loops. We then construct a weighted network model by overlaying stochastic weights based on gene expression measurements of various scRNA-seq datasets in the context of cellular differentiation and cancer. We subsequently assess global and local heterogeneity of the weighted network, namely through Forman-Ricci curvature [47, 48, 49, 50]. Further, we utilize local curvature measurements in the hypergraph model to perform pathway enrichment analysis, comparing to the graph model and a classic differential expression approach assuming gene independence. Our results indicate this approach provides a biologically meaningful measure of higher-order network heterogeneity descriptive of “stem-ness” and cancer state (globally) as well as pathway functionality (locally) in the context of cancer.

## 2 Results

Using the STRINGdb PPI database, we constructed a network model of the human interactome as a set of vertices representing unique proteins in the human proteome and a set of edges representing experimentally-determined interactions between pairs of proteins, initially resulting in a 1D graph model of the PPI network topology [14]. We then defined a higher-order PPI network model as a hypergraph based on the concept of a 2-dimensional simplicial complex [44, 46]. Using the vertices and edges of the simple graph as a starting skeleton, we built the network “up” by defining faces within the network to represent higher-order interactions between multiple proteins. Specifically, faces were identified as triplets of vertices with shared edges oriented in directions of feedback or feedforward connectivity. This 2D network model can be viewed as highly similar to the standard 1D graph model, with the same pairwise information from the graph embedded in the edges but additionally considering higher-order relationships as 2D faces (see Methods for further detail on network construction).

### 2.1 Topology of hypergraph PPI network model is distinct from graph model and exhibits higher-order organization

In examining a network model, it is reasonable to first assess the topology of the network. Topology refers to the structure of connectivity in a network; in protein-protein interaction (PPI) networks this relates to the pattern of interactions among proteins in a cell. Accordingly, we sought to examine topological properties of the two PPI network models to understand the fundamental topology that is represented by each model. We also considered an Erdős-Rényi random network of the same size to compare what would be observed in a randomly organized network [51]. Table 1 summarizes various topological parameters of the proposed 2D network model and the standard 1D graph model. The numbers of vertices |*V*| and edges |*E*| were identical in both the 1D and 2D models, due to the fact that these are defined the same in the 2D simplicial complex as in the graph. With the 2D model, the consideration of number of faces |*F*| produces a difference in the topological invariant Euler characteristic *χ* = |*V*|−|*E*| + |*F*|, indicating the two models have distinct fundamental invariant properties when considered abstractly as topological spaces. Significantly fewer 2D faces were identified in the randomly organized ER network, suggesting the PPI network exhibits increased higher-order structure compared to a random network.

**Table 1:** Topological characteristics of 1D and 2D PPI network models. 1D and 2D network models were compared in the STRINGdb PPI network (PPI) and a random Erdős-Rényi (ER) network with the same number of vertices and edges.

| Topological Measure | 1D PPI | 2D PPI | 1D ER | 2D ER |
| --- | --- | --- | --- | --- |
| Vertices (proteins) | 11 888 | 11 888 | 11 888 | 11 888 |
| Edges (interactions) | 315 130 | 315 130 | 315 130 | 315 130 |
| Faces (triplets) | 0 | 10 451 604 | 0 | 24 948 |
| Euler Characteristic ($\chi$) | -303 242 | 10 148 362 | -393 242 | -278 294 |

We then examined degree distributions in the PPI network models. Degree of connectivity was defined by a few measures: edge degree *k*_*e*_ was determined as the number of edges incident to a vertex; similarly, in the 2D model, face degree *k*_*f*_ was defined as the number of faces incident to a given vertex. Degree distributions were approximated using histogram binning. For each *k*, degree distribution *P*(*k*) was fit with a power law, *P*(*k*) ≃ *ak*^*b*^, using a linear fit on log-log transformed data, log *P*(*k*) ≃ log *a* + *b* ∗ log *k*, equivalent to the signature of a power law fit on the untransformed distribution. We report the slope coefficient *b* corresponding to the exponent of the power law and *r*^2^ of the linear fit. Both the edge-degree distribution, which was identical in the two models, along with the face-degree distribution of the 2D model revealed rough power-law behavior, which appears linear on log-log scales (Fig. 2A,B). The power law or “scalefree” property indicates a degree distribution with a high proportion of low-degree vertices and relatively few high-degree vertices [52, 53]. This property has been described in the edge degree of PPI graph models, and the scale-free property of higher-order interactions has been previously reported with regard to protein complex organization, but has yet to be explored in the context of a higher-order PPI network model [54]. Here, we also observe this property in the face-degree distribution, suggesting PPI networks organize higher-order interactions in a scale-free manner as well.

**Figure 2:**
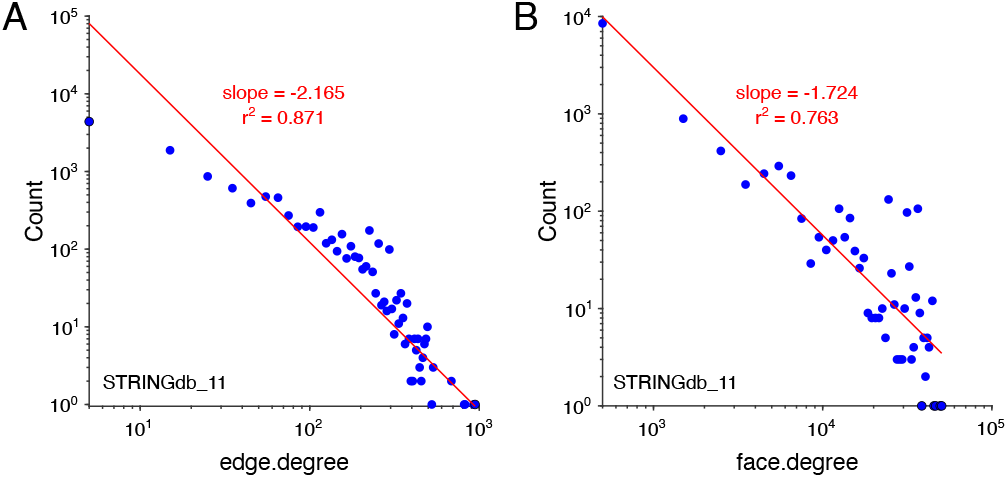
Degree distributions in PPI network. A: Log-log plot of edge-degree distribution with linear fit of power law signature. The edge-degree distribution is identical for the 1D graph and 2D simplicial complex. B: Log-log plot of face-degree distribution with linear fit of power law signature. Faces are only present in the 2D simplicial complex model.

### 2.2 Forman-Ricci curvature measures heterogeneity in weighted interaction network model

Gene expression data was overlaid onto the static PPI network topology to construct a weighted network model of stochastic protein interaction. Vertex, edge and face weights were defined using gene expression values as estimates of protein levels and principles of chemical interaction kinetics, namely the mass action law, to define interaction probabilities [55, 56, 57, 58]. Geometric weights were assigned to vertices, edges, and faces based on expression level, interaction probability and a distance-like resistance transformation (see Supplementary Methods 1 for network weighting scheme).

We then measured statistical and geometric network properties in order to assess PPI network dynamics in cells and conditions of varying gene expression (see Supplementary Methods 1 for details of datasets). With the weighted PPI network model, Forman-Ricci curvature was computed as a geometric measure of local non-uniformity in the network, allowing geometric characterization of the gene expression profile in terms of interaction network heterogeneity [42]. We also use the interaction probabilities to compute network entropy, a statistical measure of randomness and another measure of heterogeneity in the network [38].

### 2.3 Global measures of higher-order network curvature distinguish pluripotent states in differentiating stem cells

We first examine a scRNA-seq dataset of stem cell differentiation, where gene expression was measured in cells of varying degree of differentiation including pluripotent stem cells, multipotent progenitors, and differentiated cells committed to certain lineages [59]. In the weighted PPI networks, we examined global averages of Forman-Ricci curvature as well as network entropy in the 1D graph model and 2D hypergraph model. First, we observe a previously reported trend in global network entropy, wherein stem cells exhibit higher entropy that decreases with differentiation [31]; in addition, we observe a similar trend in global average Forman-Ricci curvature but in the opposite direction, wherein stem cells exhibited highly negative curvature which tended to become less negative along differentiation (Fig. 3A). In both the 1D and 2D PPI models, global average curvature was largely negative suggesting these PPI networks exhibit a generally divergent structure.

**Figure 3:**
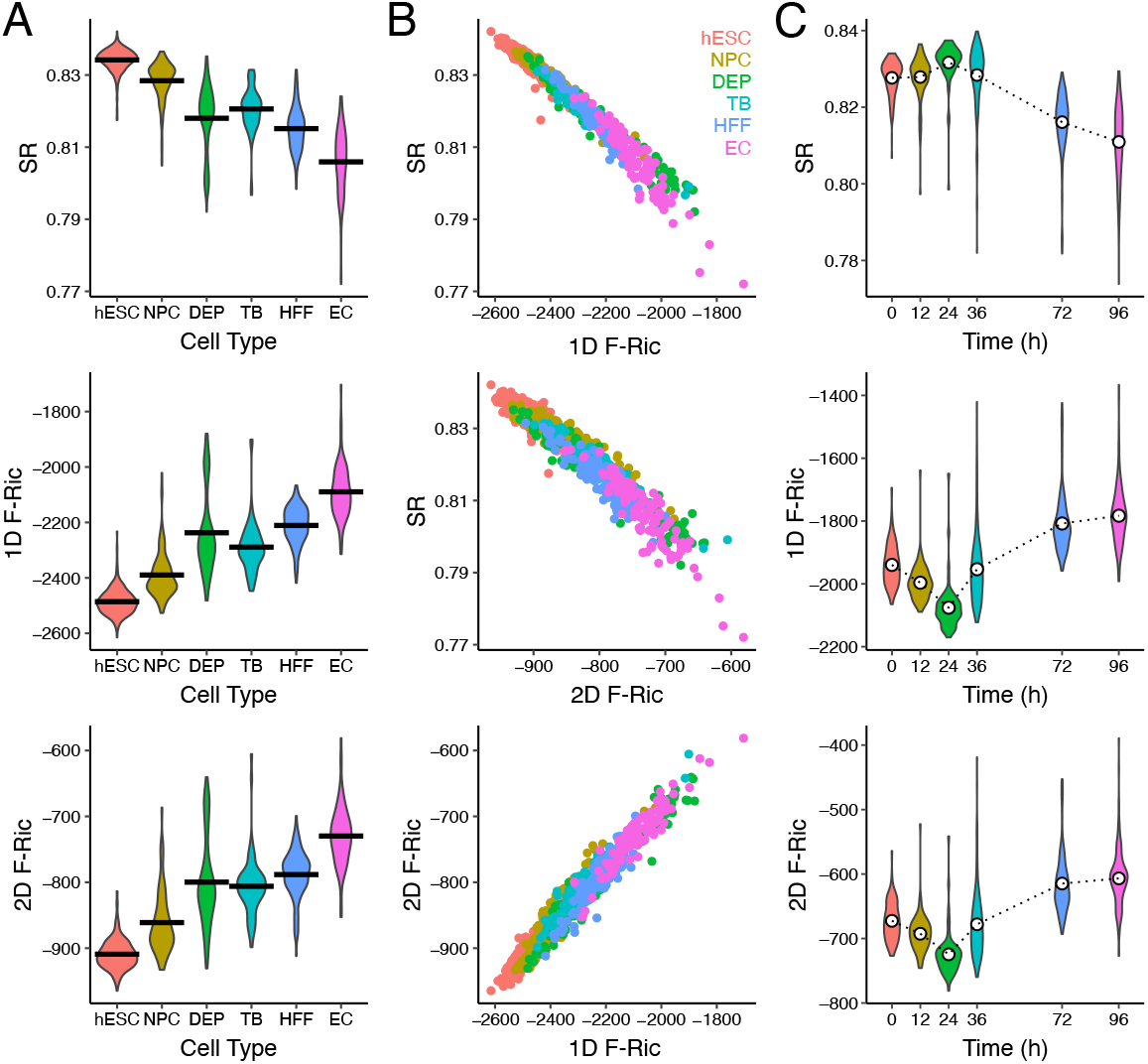
Global PPI network geometry in stem cell differentiation. A: Violin plots of PPI network global average graph entropy (SR) and Forman-Ricci curvature in the 1D graph model (1D F-Ric) and 2D simplicial complex model (2D F-Ric) across 6 increasingly differentiated cell types. B: Correlation plots of SR, 1D F-Ric and 2D F-Ric, colored by cell type. C: Violin plots of SR, 1D F-Ric, 2D F-Ric across time points after induced stem cell differentiation. Dotted lines are means at each time point.

In terms of correlation between entropy and curvature, we observed global average Forman-Ricci curvature in both models were strongly correlated with global entropy in the negative direction (1D curvature-entropy: R^2^= -0.984; 2D curvature-entropy: R^2^= -0.966); between the 1D and 2D model, global average Forman-Ricci curvature were highly correlated (1D curvature-2D curvature: R^2^= 0.980), although 2D curvature was generally shifted towards more positive values (Fig. 3B). Notably, our previous work examining curvature in the 1D graph model also found a strong correlation of Forman-Ricci curvature and entropy, but in the positive direction [42]; however, the weights of the weighted network model were previously defined using the interaction probability directly, and in this study we select a new definition of geometric weights based on the inverse of interaction probability (i.e. lower probability interactions are “longer” edges, high probability interactions are “closer”). These inverted weights are likely the reason for the discrepancy in direction, although in either case curvature and entropy were strongly correlated.

We also examined a related time-course experiment from the same stem cell dataset, where gene expression was measured as induced stem cell differentiation proceeded over time points up to 96 hours [59]. In the weighted PPI networks, global average Forman-Ricci curvature was again observed to be highly correlated in both the 1D graph and 2D hypergraph models, again with a positive shift in the 2D model (Fig. 3C). Over the time-course, curvature initially increased in magnitude (in the negative direction) up to the 24 hour time-point, and then decreased in magnitude (became more positive) at later time points. This suggests perhaps that PPI network curvature may transiently become more negative as differentiation begins but later less negative as the cell commits to a lineage.

### 2.4 Global measures of higher-order network curvature are increased in several cancer types

We examined Forman-Ricci curvature in a scRNA-seq dataset of melanoma patients including cancer cells and matching normal cells from the same patients [60]. After constructing a weighted PPI network for each cell using both the 1D graph and 2D hypergraph models, we observe on average more negative curvature globally in cancer cells relative to normal cells, which was observed across all cells in both models (Wilcoxon rank-sum test – all tumor cells vs normal cells: *p* < 0.001 in both models; Fig. 4A) and also within several individual patients (10/12) in the 1D model and all patients (12/12) in the 2D model (Fig. 4B). Again in the 2D model, curvature was generally shifted towards more positive values compared to the 1D model. The separation of cancer and normal cells suggests the magnitude of negative PPI network curvature is increased in cancer cells and could be used as a basis for classifying cancer and normal cells, for example. This distinction in network curvature globally is similar to what was observed in stem cells above, suggesting cancer cells may exhibit “stemlike” characteristics contributing to the unchecked growth and lack of differentiation observed in many cancers including melanoma. The 2D curvature appeared to show more consistent separation of the cancer and normal cells than 1D curvature or graph entropy (Supplementary Fig. S1), suggesting the 2D hypergraph model may be more informative than the 1D graph model when considering global PPI network curvature as a proxy for “stem-ness” in cancer.

**Figure 4:**
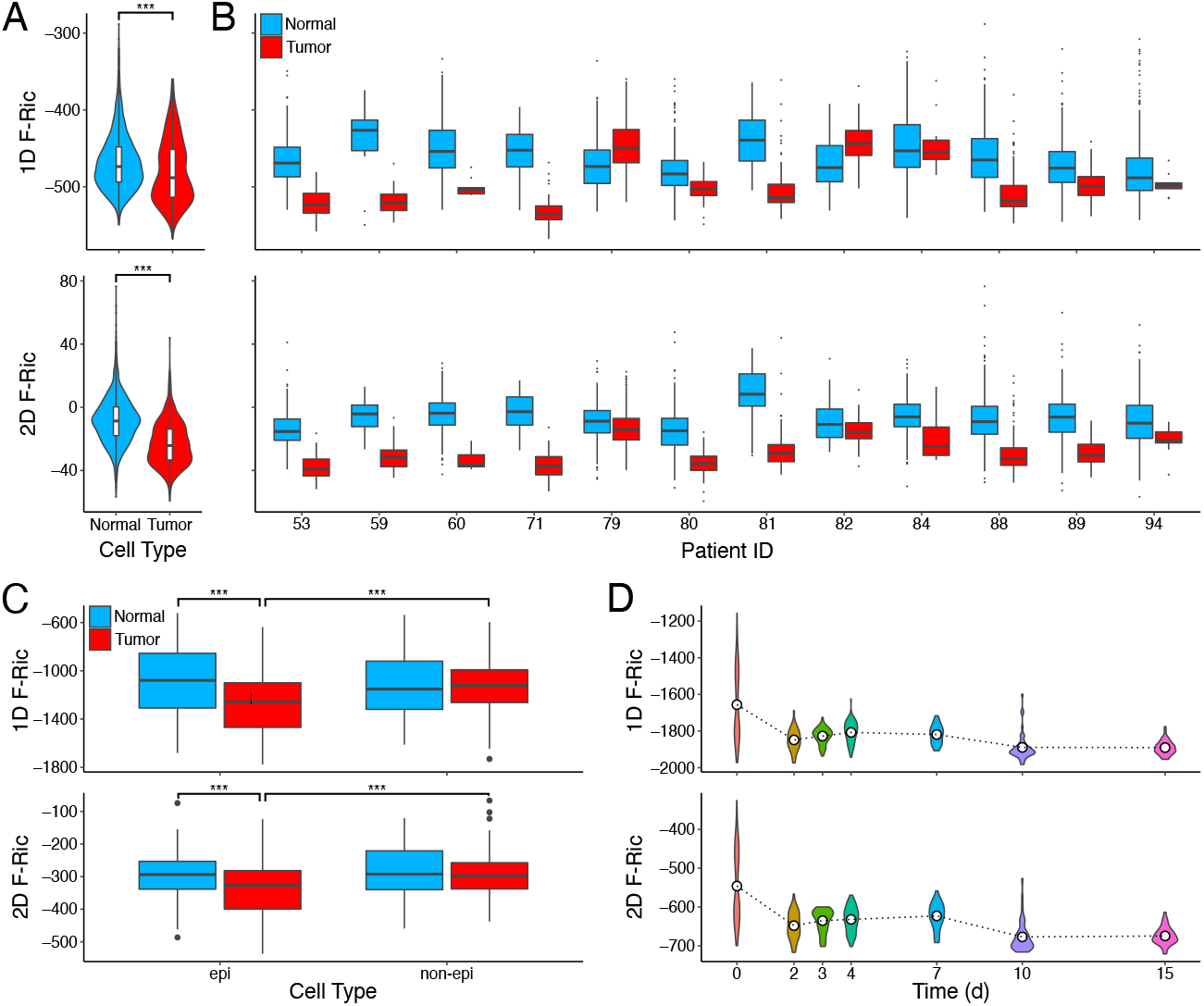
Global PPI network geometry in cancer. A: Violin plots of PPI network global average Forman-Ricci curvature in the 1D graph model (1D F-Ric) and 2D simplicial complex model (2D F-Ric) of all normal (blue) or tumor (red) cells from melanoma patients. B: Box plots of 1D F-Ric and 2D F-Ric separated by individual patients. C: Box plots of 1D F-Ric and 2D F-Ric in epithelial (epi) and non-epithelial (non-epi) cells of normal (blue) and tumor (red) samples from colorectal cancer patients. D: Box plots of 1D F-Ric and 2D F-Ric across time points after induction of EWSR1-FLI1 oncogene in A576 cells. Dotted lines are means at each time point. *** indicates *p* < 0.001.

We next examined a scRNA-seq dataset of colorectal cancer cells and matching normal cells derived from patients [61]. Examining global average Forman-Ricci curvature of the weighted PPI network models, a significant effect of tumor status and cell type on curvature were detected in both models (two-way ANOVA – 1D graph: tumor-status *p* < 0.001, cell-type *p* < 0.01, interaction *p* < 0.001; 2D hypergraph: tumor-status *p* < 0.001, cell-type *p* < 0.001, interaction *p* < 0.05). We again observed more strongly negative curvature in cancer cells compared to normal cells, as well as in epithelial cells (from which cancers arise) compared to non-epithelial stromal cells in tumors (Wilcoxon rank-sum test – normal-epithelial vs tumor-epithelial *p* < 0.001, tumor-epithelial vs tumor-nonepithelial *p* < 0.001 in both models, all other comparisons n.s.; Fig. 4C). These findings suggest the increase in negative curvature observed in cancer cells may be specific to the cancer cells themselves and is absent in the noncancerous stromal cells present in tumors.

The third cancer dataset analyzed consisted of a time-course experiment in which a Ewing sarcoma cell line was induced to express EWSR1-FLI1, a tumor-driving fusion oncogene unique to Ewing sarcoma; single cells were collected at time points after induction to measure gene expression [62]. Upon induction of the fusion gene, global average Forman-Ricci curvature was observed to initially decrease by day 2, followed by a transient increase until later time points when curvature is significantly more negative from day 0 (Fig. 4D). This trend was observed in both PPI network models. The increasingly negative curvature over time may indicate the interaction network overall becomes more divergent and thus more cancerous or “stem-like” upon expression of the EWSR1-FLI1 oncogene.

Overall, these findings demonstrate global average Forman-Ricci curvature can indicate trends of cellular pluripotency and cancer state in gene-expression-weighted PPI networks. While the 1D and 2D network models both exhibit this effect, we contend the 2D model is advantageous for its consideration of higher-order interactions not represented in the graph model and is more sensitive to biologically meaningful changes in PPI network geometry. However, these global average values are merely summary statistics for the network as a whole, which can be useful for sample-level inference but, importantly, local geometric properties contain more information for richer analysis of how individual proteins and molecular pathways contribute to the observed cell phenotype.

### 2.5 Local PPI network curvature indicates pathway functionality in cancer

We examined local curvature values in the weighted PPI networks of the melanoma dataset [60]. In order to examine changes in curvature of individual proteins, we focus on a contraction of Forman-Ricci curvature defined on vertices (Eq. 4). Vertices of the network with significantly changing curvature between the normal and cancer cells were identified. A classical differential expression approach was also used to identify up- or down-regulated genes based solely on expression, assuming gene independence. The 1D and 2D network models identified 1461/11630 (12.7%) and 1728/11630 (14.9%) genes, respectively, with significantly changing curvature in either the positive or negative direction, whereas differential expression identified only 276/11630 (2.4%) genes. We examined the number of significant genes from each method and overlap between methods and observed a high degree of overlap between the network curvature approach in the two PPI network models (Fig. 5). Of the differentially expressed genes, over half (160/276, 58.0%) of identified genes were also identified by one of the two network curvature methods, with 96 genes being determined significant by all three methods.

**Figure 5:**
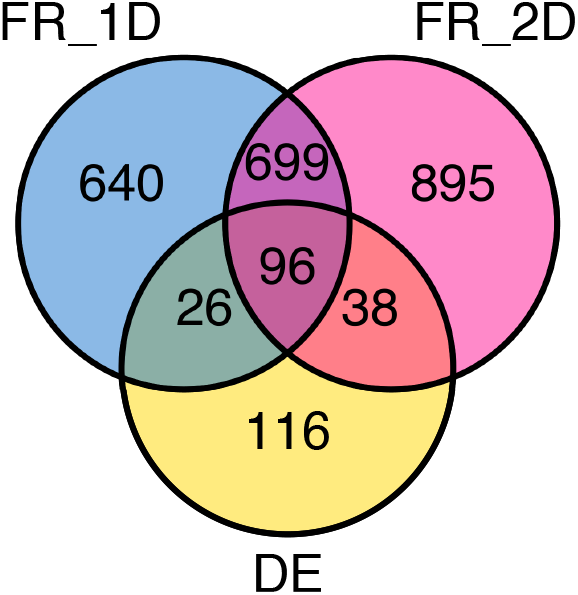
Significant gene overlaps in network curvature and differential expression. Venn diagram of the number of statistically significant genes identified by PPI network Forman-Ricci curvature in the 1D graph model (FR 1D) and 2D complex model (FR 2D) along with classic differential gene expression (DE).

We then applied a pathway analysis approach to explore molecular pathways with significantly increasing or decreasing curvature by considering genes with significantly changing vertex curvature in the 2D PPI network between normal and tumor cells. Applying Reactome pathway overrepresentation analysis on these genes [12], we found 110 pathways enriched in genes with significantly increasing curvature, along with 193 pathways with significantly decreasing curvature genes; we summarize the top 10 pathways with up or down shifts in curvature in Tables 2 and 3, respectively, with complete pathway results in Supplementary Tables 1 and 2. Because of overlapping gene sets, some pathways were redundant, such as the APC/C degradation pathway which appeared 4 times in the top 10 decreased curvature pathways. Interestingly, several of the increased curvature pathways have been previously implicated with pro-tumor involvement in melanoma, such as L1-CAM pathways and extracellular matrix organization which are involved in tumor invasiveness commonly observed in melanoma [63]. On the other end of the spectrum, several decreased curvature pathways have been implicated with tumor-suppressor involvement in melanoma and, such as PAK degradation, where PAK is a known driver of drug resistance in melanoma, and several other tumor-suppressive pathways including p53 and DNA damage response pathways (which were also significantly enriched but not shown in top 10, see Supplementary Table 2) [64, 65]. Because of the relationship of curvature and robustness, these findings suggest oncogenic pathways with increasing curvature also increase in robustness in the cancer cells, while tumor suppressive pathways correspondingly decrease in robustness, thereby contributing to the cancer phenotype. Therefore, estimating robustness through PPI network curvature may be useful to indicate gene and pathway functionality in cellular processes such as cancer development.

**Table 2:** Increased PPI Curvature Pathways in Melanoma. Using vertex curvature of the 2D PPI network model, genes with significantly increasing curvature values compared between normal and tumor cells were fed into Reactome Pathway Analysis. Top 10 enriched pathways are summarized by pathway name, number of significant genes, false-determination rate (FDR), and average shift in curvature (ΔRic_2D_).

| Pathway Name | #gene | FDR | $\Delta\text{Ric}_{2D}$ |
| --- | --- | --- | --- |
| Regulation of HSF1-mediated heat shock response | 22 | 1.36e-4 | +51.70 |
| Interaction between L1 and Ankyrins | 12 | 1.36e-4 | +47.27 |
| Post-translational protein phosphorylation | 20 | 3.52e-4 | +125.63 |
| Attenuation phase | 13 | 3.52e-4 | +63.82 |
| Processing of DNA double-strand break ends | 18 | 3.52e-4 | +169.67 |
| Cellular response to heat stress | 22 | 3.59e-4 | +51.69 |
| L1CAM interactions | 21 | 6.13e-4 | +56.87 |
| Elastic fibre formation | 12 | 6.28e-4 | +42.76 |
| G2/M DNA damage checkpoint | 16 | 6.28e-4 | +183.55 |
| Extracellular matrix organization | 37 | 7.67e-4 | +58.11 |

**Table 3:** Decreased PPI Curvature Pathways in Melanoma. Using vertex curvature of the 2D PPI network model, genes with significantly decreasing curvature values compared between normal and tumor cells were fed into Reactome Pathway Analysis. Top 10 enriched pathways are summarized by pathway name, number of significant genes, false-determination rate (FDR), and average shift in curvature (ΔRic_2D_).

| Pathway Name | #gene | FDR | $\Delta Ric_{2D}$ |
| --- | --- | --- | --- |
| APC/C:Cdc20 mediated degradation of Securin | 34 | 7.65e-13 | -340.13 |
| The role of GTSE1 in G2/M progression after G2 checkpoint | 36 | 2.58e-12 | -284.47 |
| APC/C:Cdh1 mediated degradation of Cdc20 and other APC/C:Cdh1 targeted proteins in late mitosis/early G1 | 34 | 2.58e-12 | -340.13 |
| NIK→ noncanonical NF-kB signaling | 31 | 2.58e-12 | -293.12 |
| APC/C:Cdc20 mediated degradation of mitotic proteins | 34 | 2.98e-12 | -340.13 |
| Regulation of activated PAK-2p34 by proteasome mediated degradation | 28 | 2.98e-12 | -305.36 |
| Activation of APC/C and APC/C:Cdc20 mediated degradation of mitotic proteins | 34 | 2.98e-12 | -340.13 |
| FBXL7 down-regulates AURKA during mitotic entry and in early mitosis | 29 | 2.98e-12 | -304.05 |
| SCF-beta-TrCP mediated degradation of Emi1 | 29 | 2.98e-12 | -304.05 |
| Regulation of ornithine decarboxylase (ODC) | 28 | 2.98e-12 | -305.36 |

## 3 Discussion

Because cells are inherently dynamic systems formed of molecular interactions, effective models of the interaction dynamics within cells can reveal how the “interactome” varies in cells of differing phenotypes, including pathological (i.e. cancer) and healthy physiological states. In this study, we modeled interactome dynamics by incorporating gene expression information into a weighted protein-protein interaction (PPI) network and measured network heterogeneity by Forman-Ricci curvature. Heterogeneity in interaction dynamics can lend to increased robustness [43]. For example, a given cancer cell may be difficult to treat if multiple, diverse growth-stimulating pathways drive the cancer’s growth; alternatively, a cancer that depends on a single oncogenic pathway may be easy to treat with targeted therapy.

Critically, the most common model of PPI networks, the graph, is arguably limited insofar that a graph only considers pairwise relationships between two proteins at most. Protein interactions are not strictly pairwise and can involve multiple proteins that coordinate in interactions and molecular processes. We sought to develop a model of higher-order structure in the interactome to address the limitation of the graph model to pairwise information. Here, we introduced a hypergraph model of the PPI network based on a 2-dimensional (2D) simplicial complex that represents feedforward and feedback structures in the network as 2D triangular faces. Importantly, we limited our examination to triangles, which correspond to feedback or feedforward loops involving three proteins, but the approach outlined can easily be extended to represent higher-order interactions of more proteins as polygonal faces with more sides (quadrangles, etc.).

We first examined topological properties of this 2D model and drew comparisons between the standard 1-dimensional (1D) graph model, as the two models represent much of the same protein interaction information but are fundamentally distinct. We contend the standard graph model of PPI networks is essentially a “skeleton” of pairwise interactions in the interactome and that by considering higher-order relationships in the structure of the network, we approach a more comprehensive model of the system of diverse protein interactions within cells. The 2D faces of our model then serve as important representations of higher-order relationships among interacting proteins that contribute to molecular pathways and dynamic cellular processes.

Next, we applied the higher-order network model to analyze publicly available gene expression (RNA-sequencing) datasets in the context of cellular differentiation and cancer experiments. We constructed a weighted interaction network model, assigning geometric weights reflecting stochastic interaction rates to each of the vertices, edges, and faces in the case of the 2D simplicial complex model. We then measured geometric properties of the weighted network, namely Forman-Ricci curvature, as a means to quantify heterogeneity in the network. This heterogeneity can serve as a measure of network dynamics, i.e. robustness, to describe the stability or fragility of the network on a global (sample-level) and local (protein- or pathway-level) scale. Importantly, the 2D network model is suitable for geometric analysis by incorporating higher-dimensional information, which is in line with the complete definition of Forman-Ricci curvature [44, 47].

We examined Forman-Ricci curvature of the 2D weighted PPI network on a global scale and compared with results of the 1D graph model, finding network curvature in both models was highly correlated and negative on average, although with a shift towards less negative curvature overall in the 2D model. The global average curvature in either model distinguished undifferentiated stem cells from differentiated cell types, as well as cancer cells from normal cells; curvature in the 2D model, however, appeared to provide a more consistent separation, suggesting the higher-order model may be a more accurate representation of the PPI network.

To further demonstrate the biological relevance of the proposed higher-order model and network geometric approach, we applied pathway analysis using local curvature to identify proteins with changes in curvature between patient-matched cancer and normal cells. Despite the overall shift towards negative curvature in cancer, we found that proteins with increased curvature appeared to enrich for several pro-cancer pathways, along with decreased curvature in tumor-suppressive pathways, suggesting a relationship of the measured local curvature with the robustness of these pathways which may be related to their functionality in the cancer cells. We conclude this model provides a feasible approach to analyzing how higher-order relationships in the interactome influence cellular phenotype and function, especially in terms of quantifying stability and fragility in the network which may be a valuable tool to identify potential therapeutic strategies in cancer, for example.

Future directions of this research include further exploration of higher-order models of PPI networks and applying the model to investigate biological questions about how the interactome governs cellular phenotype and behavior. The geometric framework that we apply to analyze PPI networks allows for application of additional geometric tools, such as geometric flows (i.e. discrete Ricci flow) that can be used for change detection and prediction of network dynamics [49, 66, 67]. Fortunately, many of the same statistical and geometric network properties measured in graphs can be considered in generalized network models through extensions of definitions including entropy and Forman-Ricci curvature [44, 68, 69, 70]. Our characterization of the proposed model, while intended to be thorough, is by no means an exhaustive analysis and is meant to illustrate a higher-order PPI network model and a network geometric approach to studying interaction dynamics.

## 4 Methods

### 4.1 Higher-order protein interaction network model

A graph *G* = (*V, E*) was defined as the sets of vertices *V* and edges *E* corresponding to proteins and pairwise interactions, respectively. Interactions were defined based on the STRINGdb PPI database, including an experimental confidence score cutoff. To represent higher-order relationships involving more than 2 proteins, we defined a hypergraph based on the concept of a 2-dimensional simplicial complex, where *simplicial* complex indicates that all faces are 2-simplices, or triangles. Similar to the definition of the graph, we defined a 2-dimensional complex *C* as a set of vertices *V*, edges *E* and faces *F*, notated as *C* = (*V, E, F*). Faces were identified as all triplets of vertices with edges among all vertices arranged in feedforward (+) or feedback (–) orientation (Fig. 6). The higher-dimensional structure therein serves to represent higher-order relationships among interacting proteins while still incorporating pairwise interactions from the (1-dimensional) graph (see Supplementary Methods 1 for additional details on network construction).

**Figure 6:**
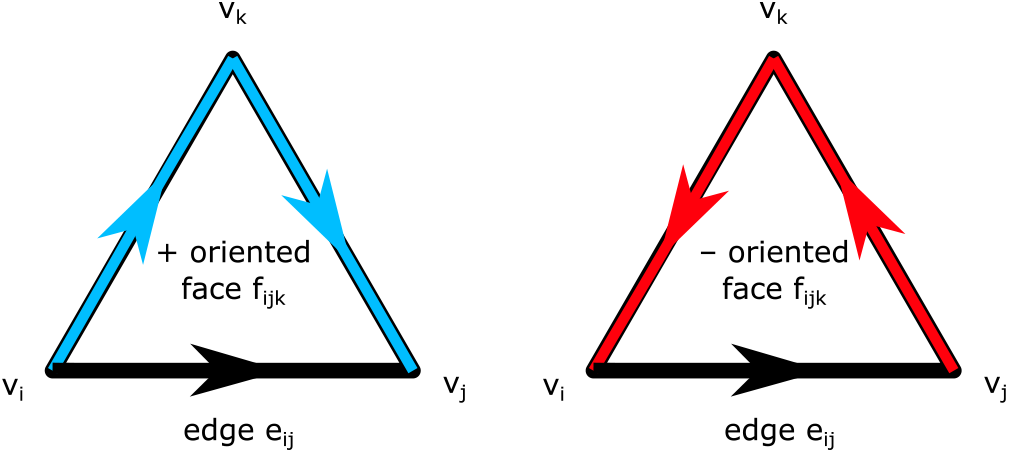
Directed network face orientation. Face orientation with respect to a given edge is determined by the direction of the corresponding edges in the face. In the same direction is positive (+) oriented, or feedforward; in the opposite direction is negative (–) oriented, or feedback.

### 4.2 Weighted protein interaction network model

We consider the PPI network as an underlying structure and incorporate measured data about the system (i.e. gene expression) to construct a weighted network model. In the PPI network, gene expression information (i.e. RNA sequencing) provides an estimate of protein levels corresponding to the vertices of the interaction network [55]. This weighted PPI network is then an effective model of cellular dynamics, allowing assessment of how protein levels and interaction rates vary over a time-course or across differing cell types. We selected a stochastic weighting scheme which relates protein levels to interaction probability by the mass-action principle (see Supplementary Methods 1 for weighting scheme). Network weights were defined for each sample (i.e. single-cell) of gene expression data, allowing measurement of local and global network properties for each sample.

### 4.3 Discretization of Ricci curvature

Ricci curvature, a geometric measure of deviation from “flatness”, can be used to quantify local non-uniformity. We compute network Ricci curvature as a means to quantify heterogeneity in the weighted PPI network model. Importantly, network models are inherently discrete, therefore a discretization of Ricci curvature is required to consider this geometric property in a network model. A few discrete definitions exist for Ricci curvature on networks [50, 71, 72]; here we focus on Forman-Ricci curvature because of its unique amenability to extension to the higher-dimensional network model [44, 47]. Forman-Ricci curvature is derived through a combinatorial approach which applies to the general class of CW-complexes including graphs and simplicial complexes. In this sense, a graph is considered a complex of vertices (0-cells) and edges (1-cells) glued together at their boundaries, i.e. their vertices. This notion can be extended to hypergraphs through the 2-dimensional simplicial complex model, by considering also faces (2-cells) glued together at boundary edges and vertices. Then, Forman’s approach derives a combinatorial formula for discrete Ricci curvature which depends only on the weights of the edge, adjacent vertices and faces, and parallel edges.

In brief, the derivation of Forman-Ricci curvature is based on a combinatorial ana-log of Bochner-Weitzenboöck decomposition of the Riemannian Laplacian operator:

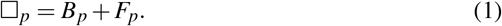

Essentially, the Hodge Laplacian □_*p*_ can be decomposed into a “rough” Laplacian *B*_*p*_ and a curvature term *F*_*p*_ capturing how the two Laplacians differ. Because the Laplacian relates to diffusion on a manifold, curvature on a weighted PPI network can then quantify how diffusion (i.e. information flow through protein interactions) differs from expected in an un-curved (flat) network.

Forman’s approach yields the following explicit formula for discrete Ricci curvature on a weighted complex with edges *e*, vertices *v* and faces *f* :

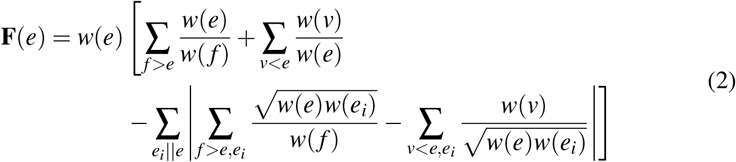

where *f* > *e* implies *f* is a face of *e* and *v* < *e* implies *v* is a vertex of *e*, and *e*_*i*_∥*e* implies *e*_*i*_ is a *parallel* edge of *e* meaning it shares a face or a vertex but not both.

The proposed 2-dimensional complex model of PPI networks can be analyzed geometrically by properly defining face weights (see Supplementary Methods 1 for weighting scheme) and applying formulae which account for the 2D faces. Because Forman-Ricci curvature is defined for complexes of any dimension, the definition Eq. 2 is readily applicable to the proposed model. In fact, the original derivation of Forman-Ricci curvature directly considers 2-dimensional faces; applications on 1-dimensional graphs consider a simplified definition that disregards all face-related terms [47, 48, 50]:

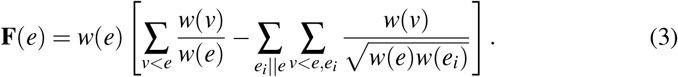

While Forman-Ricci curvature is defined on *edges*, it can be contracted to a vertex curvature **F**(*v*) and then a global average of curvature **F**_*GA*_ can be computed using the stationary distribution *π* of the network, where *π*_*i*_ designates the equilibrium probability of a Markov random walk at each vertex:

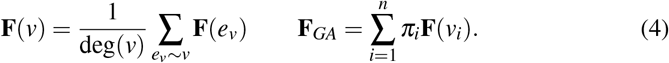

## Acknowledgements

This work was supported by the National Science Foundation [ECCS-1749937 to K.A.M. and R.S.]; the U.S. Air Force Office of Scientific Research [FA9550-18-1-0130 to K.A.M. and R.S.]; and the German-Israeli Foundation [I-1514-304.6/2019 to E.S.].

## Author Contributions

All authors contributed to the intellectual design of the study. K.A.M. developed computational code, identified relevant datasets, and analyzed results. E.S. and R.S. provided additional guidance throughout the study. All authors contributed to scientific discussion and preparation of the manuscript.

