## Supplementary Methods 1 for "Beyond Pairwise Interactions: Higher-Order Dynamics in Protein Interaction Networks"

### Construction of PPI Network Topology

Protein-protein interaction data was integrated from STRINGdb version 11[1]. Interaction data was provided as a list of edges, with start- and end-vertices corresponding to individual proteins of each interaction. To ensure high-confidence interactions, an experiment score cut-off of >0.9 was applied. Additionally, a previously described PPI sparsification procedure was applied to remove likely false positive interactions based on GO subcellular localization terms[2].

*1-Dimensional graph model*

To represent the PPI network as a (1-dimensional) graph, an adjacency matrix of size $|V|\times|V|$ was defined with entries of 1 or 0 corresponding to the presence of an edge between vertices. After taking the largest strongly connected component, this graph served as a the topological basis for the 1-dimensional (1D) interaction network model.

*2-Dimensional simplicial complex (hypergraph) model*

Extension of the 1D graph to a hypergraph was accomplished by defining a 2-dimensional simplicial complex with 2-dimensional faces where triangles were formed by three vertices sharing three respective edges. Each triangle was identified by its unique triplet of vertices. The resulting 2-dimensional simplicial complex was used as the topological basis for the 2-dimensional (2D) interaction network model.

### Weighted PPI Network Model

The weighted network was defined on the intersection of the gene expression data and the PPI topology, that is vertices without matching gene expression data or genes without corresponding vertices in the PPI network were discarded. From the resulting network containing the intersection of the data, the largest strongly connected component was taken as the graph model. In the 2D model, faces were identified as described above. Weights were then assigned to the vertices, edges, and faces if present.

*Stochastic interaction network weights*

For a given sample of gene expression, vertex weights were defined as the (pre-processed) expression value $E_i$ of the gene corresponding to vertex $i$ in the PPI network as a direct estimate of protein level[3]:

$$w(v_i) = E_i.$$

Interaction probabilities were defined (as previously described[4]) using the mass action law, which states the rate of interaction is proportional to the amount of the reactant proteins, corresponding to the product of the expression values of the two vertices of a given edge. These mass-action products were then normalized by the sum of all outgoing products at each vertex. The normalized mass-action products $p_{ij}$ represent the probability of interaction of a given protein with each of its interacting partners, similar to the transition probability of a Markov random walk:

$$p_{ij} = \frac{E_i E_j}{\sum_{k \in N(i)} E_i E_k},$$

where $N(i)$ denotes the indices of neighboring vertices adjacent to vertex $i$.

We then utilized the interaction probability to define geometric weights on edges using a resistance transformation, defined as the inverse of the product of the out-degree $deg_{out}$ of the start vertex times the interaction probability $p_{ij}$ of that edge:

$$w(e_{ij}) = \left(deg_{out}(v_i) p_{ij}\right)^{-1}.$$

This resistance transformation directly relates to commuting times of a random walk and satisfies metric properties except for the symmetry condition[5,6]. Intuitively, a low interaction probability equates to a large resistance and vice versa, thereby ascribing a geometric ``distance" from interaction probability which we consider as edge weights. By using this definition, the network inherently becomes a directed network (even if the topology was originally undirected) because the probability of interaction and resulting resistance metric between two proteins is generally not equal in both directions.

In the 2D (hypergraph) model, faces were defined as all sets of 3 vertices with shared edges among all vertices. In the directed network, faces are only considered in the feedforward (+) or feedback (–) orientations with respect to a given edge (Fig. 2 of main text). Face weights were defined directly from the edge weights of a face as the geometric mean of the edge weights squared, analogous to area as the square of length. Because of face orientation, weights of edges in the relevant orientation were considered and the orientation was used to ascribed a positive or negative sign to the face weight:

$$w\left(f_{ijk}^{+}\right) = \left(w(e_{ij})w(e_{ik})w(e_{kj})\right)^{2/3}, \quad w\left(f_{ijk}^{-}\right) = -\left(w(e_{ij})w(e_{jk})w(e_{ki})\right)^{2/3}.$$

*Network Forman-Ricci curvature computation*

Eqs. (2) and (3) of main text were applied as the definitions of Forman-Ricci curvature on the 2D and 1D models, respectively. These combinatorial definitions, which are used to calculate curvature on *edges*, depend solely on the weights of the edge of interest, incident vertices (and faces in the 2D model), and parallel edges. The resulting edge curvature values were contracted to vertex curvature as the mean of incoming minus outgoing edge curvatures, and a global weighted average of vertex curvature was calculated using the stationary (invariant) distribution $\pi$ of the network as the contribution of each vertex in the weighted average (Eq. (4) of main text). $\pi$ was computed as the normalized first eigenvector of the interaction probability matrix $p_{ij}$, representing the equilibrium distribution of a Markov random walk over the network. Altogether, for each sample of gene expression data this approach resulted in a length |E| vector of edge Forman-Ricci curvatures, a length |V| vector of vertex curvatures and a single scalar global weighted average curvature.

*Network entropy computation*

Network entropy was computed using the same interaction probabilities $p_{ij}$ described above[4]. From this Markov-chain-like stochastic network, Shannon entropy $S_i$ was computed locally at each vertex; local entropy was then averaged over the stationary distribution $\pi$ of the network to compute a global entropy rate $SR$:

$$S_i = \sum_{j \in N(i)} p_{ij} \log p_{ij}\,, \qquad SR = \sum_{i=1}^{n} \pi_i S_i,$$

where stationary distribution $\pi$ was calculated as above.

Shannon entropy measures the expected information content of a probabilistic event; in a PPI network this can be interpreted as the relative randomness of a protein's interactions with its partners. The maximum possible entropy at a vertex occurs when all neighboring edges have equal probability of interaction and the minimum occurs when all probability lies on a single edge. In this sense, higher entropy describes a less ``committed'' state of the network and decreases as the network ``commits'' to specific proteins and interactions[7].

**Datasets and computation**

Publicly available single-cell RNA-sequencing (scRNA-seq) datasets were downloaded from NCBI GEO database, accession codes: GSE75748, GSE72056, GSE81861, GSE130019. Pre-processing of raw RNA read counts included quantile normalization, in order to normalize relative counts between samples, and $\log_2$-transformation to reduce the influence of highly expressed genes.

*GSE75748: Stem cell differentiation*

This dataset contains two scRNA-seq experiments examining human stem cell differentiation[8]. The first experiment is a cell-type experiment consisting of n=1,018 single cells derived from human embryonic stem cells and identified as one of 6 cell types: *hESC* human embryonic stem cell, *NPC* neuronal progenitor cell, *DEP* definitive endoderm progenitor, *TB* trophoblast-like cell, *HFF* human foreskin fibroblast, *EC* endothelial cell. The second experiment is a time-course experiment consisting of n=758 single cells induced to differentiate from H9 stem cells and collected at specific time intervals: 0h, 12h, 36h, 72h, 96h.

*GSE72056: Melanoma*

This dataset contains scRNA-seq data of melanoma patient samples including tumor cells and matched normal cells[9]. The data contains a total of n=4,513 single cells from 19 patients, including $n_n$=3,256 normal cells and $n_t$=1,257 tumor cells. To compare within or between patients, a filter was applied to remove patients with less than 5 normal or 5 tumor samples, leaving 12 patients and n=3,424 cells total ($n_n$=2294, $n_t$=1,130).

*GSE81861: Colorectal cancer*

This dataset contains scRNA-seq data of colorectal cancer patient samples including tumor cells and matching normal cells[10]. The data contains a total of n=641 single cells, including $n_n$=266 normal cells and $n_t$=375 tumor cells. Each cell was labelled as one of 7 cell types: epithelial, fibroblast, endothelial, Bcell, Tcell, macrophage, mast cell. Any cell with no labelled cell type was discarded. Additionally, any cell types with less than 5 normal or 5 tumor samples were removed from analysis. For comparison to epithelial cells, the primary cell type in intestinal mucosa from which colorectal adenocarcinoma arises ($n_e$=475), all non-epithelial cells ($n_{ne}$=145) were combined into a single group.

*GSE130019: Ewing sarcoma cell line*

This dataset contains a scRNA-seq experiment examining a Ewing sarcoma cell line[11]. A576 cell lines with EWS1-FLI1 fusion mutation, an oncogene specific to Ewing sarcoma, were treated with DOX to suppress the expression of the fusion gene for 7 days, then were released from DOX treatment allowing re-expression of the oncogene. Cells (n=598) were collected at time points after DOX cessation: 0d, 2d, 3d, 4d, 7d, 10d, 15d.

*Pathway enrichment analysis*

To determine the biological function of genes corresponding to proteins of interest in the PPI network, we apply a pathway enrichment analysis that statistically associates a set of genes with molecular pathways of known biological function. Pathway enrichment analysis was performed by first performing a non-parametric Wilcoxon rank-sum test on the vertex curvature of each gene between conditions (i.e. normal cells vs. tumor cells) to compute the ratio of mean vertex curvature $\Delta$Ric and significance *p*-value between the conditions. False discovery rate (FDR) was adjusted from the *p*-value using the Benjamini-Hochberg step-up procedure[12]. Selection cutoffs of FDR<0.05 and absolute $\Delta$Ric>2 were applied to select genes with significant effect size, which were further split into increasing or decreasing curvature genes. Then, gene symbols were directly fed into the Reactome online pathway analysis tool, which determines pathway overrepresentation using a hypergeometric distribution[13]. Enriched pathways were selected based on FDR<0.05 and >10 genes in the pathway.

*Code and computation*

Data processing and analysis code were written in MATLAB and R. For efficient computation of large gene expression datasets, a parallelized algorithm was employed on a high-performance compute cluster, with the full algorithm running in approximately 6 seconds per sample (i.e. single-cell) of data. All relevant code and data is available upon request to the authors.
