## Supplementary figures and images for "Beyond Pairwise Interactions: Higher-Order Dynamics in Protein Interaction Networks"

### Supplementary Figure 1

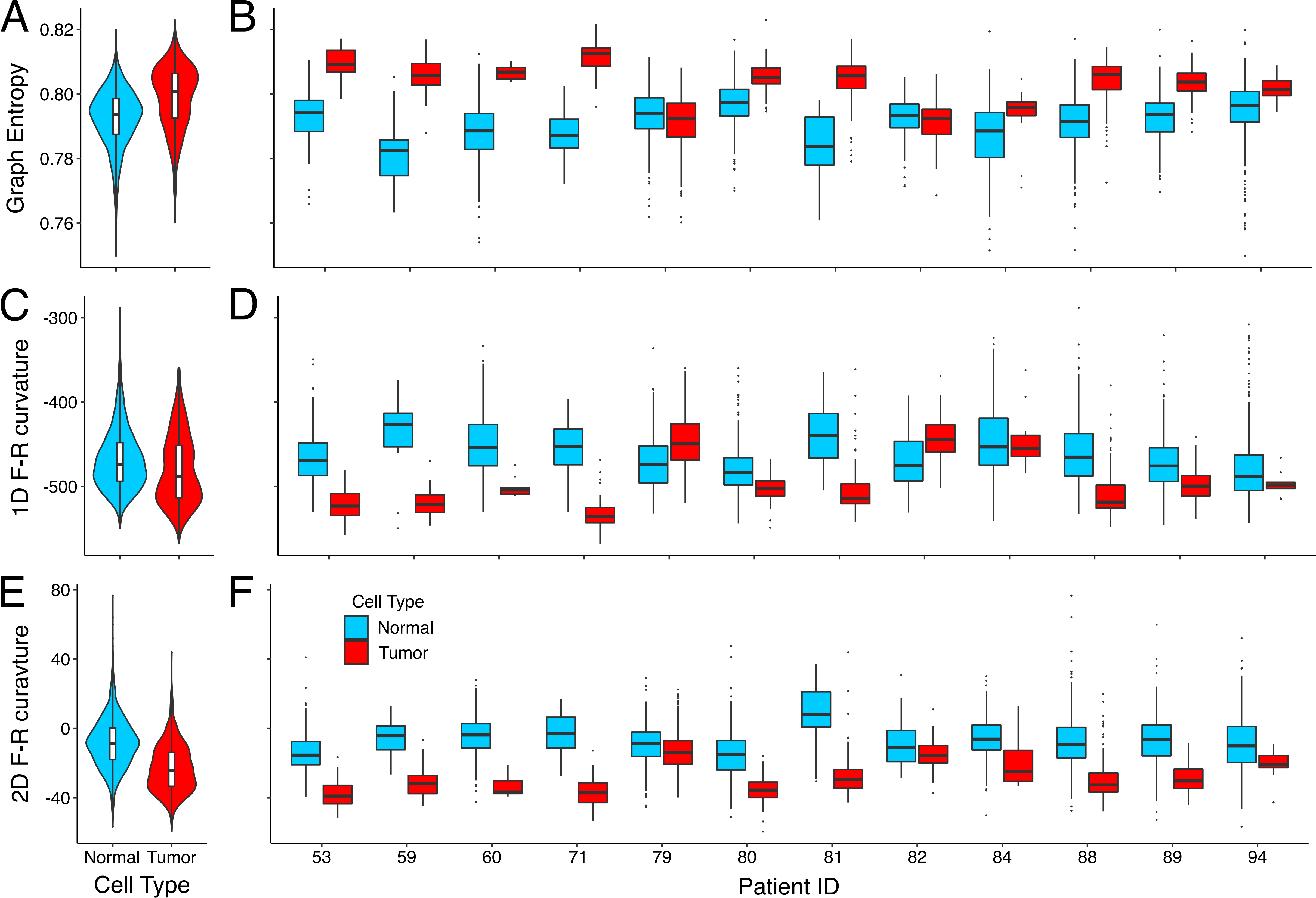
